# Genome-wide Association Study Of Plasma Proteins Identifies Putatively Causal Genes, Proteins, And Pathways For Cardiovascular Disease

**DOI:** 10.1101/136523

**Authors:** Chen Yao, George Chen, Ci Song, Michael Mendelson, Tianxiao Huan, Annika Laser, Hongsheng Wu, Jennifer E. Ho, Paul Courchesne, Asya Lyass, Martin G. Larson, Christian Gieger, Johannes Graumann, Andrew D. Johnson, Shih-Jen Hwang, Chunyu Liu, Karsten Suhre, Daniel Levy

## Abstract

Identifying genetic variants associated with circulating protein concentrations (pQTLs) and integrating them with variants from genome-wide association studies (GWAS) may illuminate the proteome’s causal role in disease and bridge a GWAS knowledge gap for hitherto unexplained SNP-disease associations. We conducted GWAS of 71 high-value proteins for cardiovascular disease in 6,861 Framingham Heart Study participants followed by external replication. We comprehensively mapped thousands of pQTLs, including functional annotations and clinical-trait associations, and created an integrated plasma-protein-QTL searchable database. We next identified 15 proteins with pQTLs coinciding with coronary heart disease (CHD)-related variants from GWAS or tested causal for CHD by Mendelian randomization; most of these proteins were associated with new-onset cardiovascular disease events in Framingham participants with long-term follow-up. Identifying pQTLs and integrating them with GWAS results yields insights into genes, proteins, and pathways that may be causally associated with disease and can serve as therapeutic targets for treatment and prevention.

## Introduction

Considerable progress has been made in identifying genetic underpinnings of coronary heart disease (CHD),^1-4^ which remains the leading cause of death worldwide.^5^ Proteins are the functional products of the genome and serve as critical factors for biological processes involved in health and disease as well as primary drug targets. Numerous proteins have been reported to be associated with CHD; it is often difficult, however, to establish with certainty whether CHD-associated proteins are causally related to risk or simply represent downstream markers of disease-related processes. Identifying genetic variants associated with protein levels (protein quantitative trait loci; pQTLs), characterizing pQTLs that also are associated with CHD from genome-wide association studies (GWAS), and inferring causality may provide novel insights into the roles of genetic variants, genes, and the proteins they code in the pathogenesis of CHD. To date, most pQTL studies^6-15^ have been based on small sample sizes or did not conduct prospective testing of associations between protein levels and clinical disease.

To address a GWAS knowledge gap for genetic variants of unknown relevance to CHD, we conducted a multistage study (Figure 1) consisting of GWAS of high-value cardiovascular disease (CVD) plasma proteins that were measured in Framingham Heart Study (FHS) participants, followed by external replication in participants from the Cooperative Health Research in the Region of Augsburg (KORA) F4 study^12^ and from other protein GWAS. We integrated pQTLs with genetic variants from CHD GWAS databases^1-4^ and employed Mendelian randomization (MR)^16^ to reveal proteins with potentially causal effects on CHD. Last, we tested proteins for association with new-onset CHD events in FHS participants with long-term follow-up. We hypothesized that a strategy of protein GWAS followed by causal testing and prospective association with CHD outcomes would identify putatively causal genes, proteins, and pathways for CHD and highlight novel targets for its prevention and treatment.

**Figure 1.**
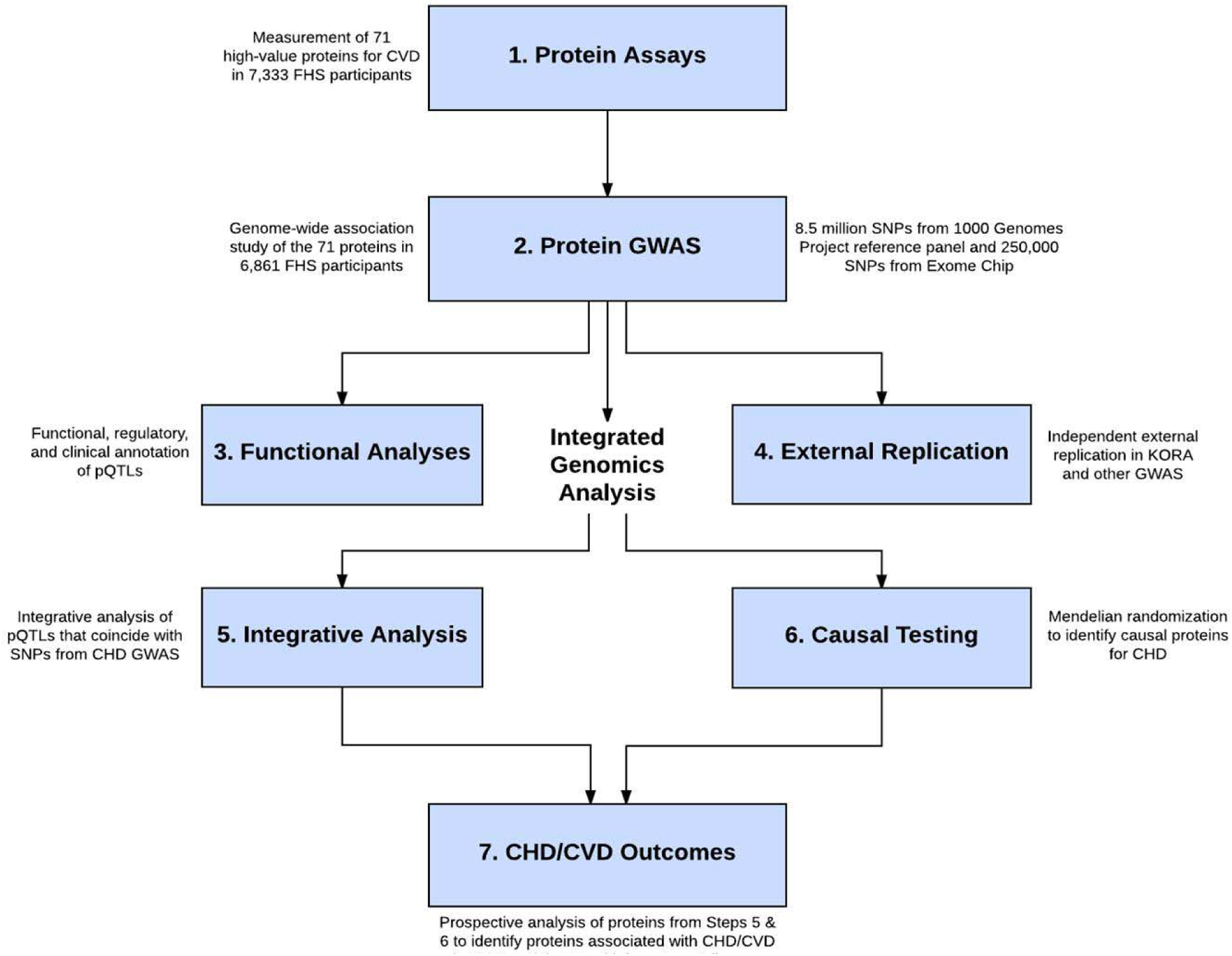
Flowchart of Study Design. The study consisted of seven steps: 1) selection and measurement of 71 high-value plasma proteins for atherosclerotic CVD via multiplex immunoassays in 7,333 FHS participants, 2) GWAS of the 71 proteins in 6,861 FHS participants to identify genome-wide significant pQTLs, 3) functional enrichment analyses of the identified pQTLs, 4) independent external replication of the sentinel pQTLs in KORA and other previous GWAS, 5) integrated analysis to pQTLs that coincide with CHD SNPs from GWAS, 6) identification of causal proteins for CHD using a Mendelian randomization approach, 7) association analysis of proteins from steps 5 and 6 with risk for incident CHD death and CVD death in 3,520 FHS participants age 50 years or older with available long-term follow-up. Abbreviations: CHD = coronary heart disease; CVD = cardiovascular disease; FHS = Framingham Heart Study; GWAS = genome-wide association study; KORA = Cooperative Health Research in the Region of Augsburg Study; pQTL = protein quantitative trait locus (i.e. genetic variant associated with protein level); SNP = single nucleotide polymorphism

## Findings

### Discovery Set

Seventy-one proteins, selected *a priori* based on prior evidence of association with CVD, were measured in 7,333 FHS participants (Table S1). The sample size available for GWAS was up to 6,861 participants (mean age 50 years, 53% women); clinical characteristics of the discovery sample are summarized in Table S2.

### pQTL Mapping

With a GWAS sample size of ∼6,800 participants and a significance threshold of p< 5x10^−8^, our study had 80% power to detect a pQTL that explained ≥0.6% of variance in protein levels (Table S3). We identified 1,793 insertion/deletion polymorphisms for 57 proteins (Table S4) and 20,495 pQTLs with Reference SNP cluster IDs for 60 proteins (Table S5), including 11,974 cis-pQTLs representing 39 sentinel *cis*-pQTL loci for 39 proteins (Figure 2a; Table S6) and 8,521 trans-pQTLs representing 91 sentinel *trans*-pQTL loci for 48 proteins (Figure 2b; Table S6). Pruning the 1000 Genomes Project (1000G) reference panel^17^ GWAS pQTLs (linkage disequilibrium r^2^< 0.2) yielded 4,588 non-redundant variants (Table S7). Thirty-four pQTLs were rare variants (minor allele frequency< 1%) associated with 18 proteins (Table S8).

**Figure 2.**
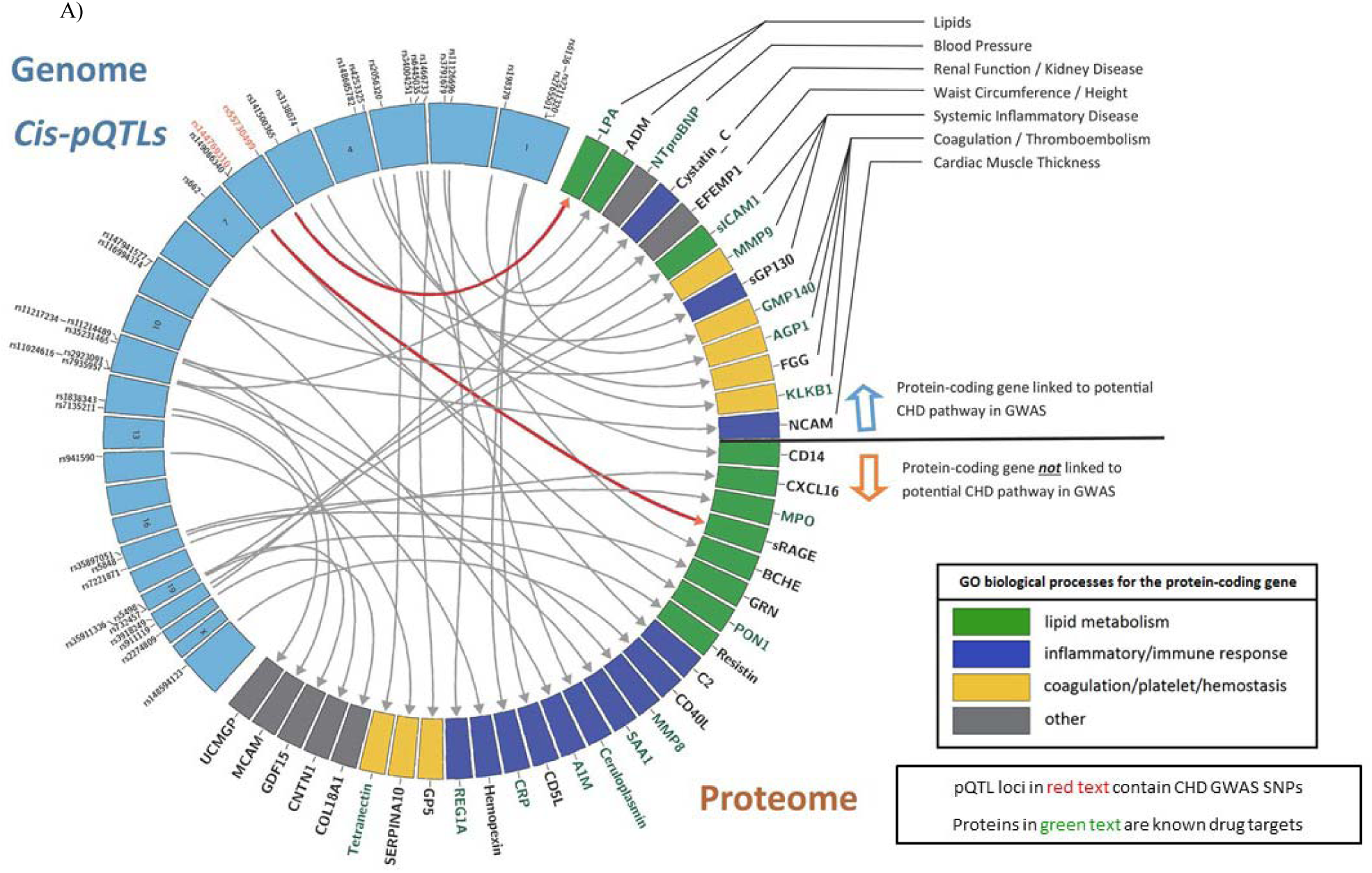
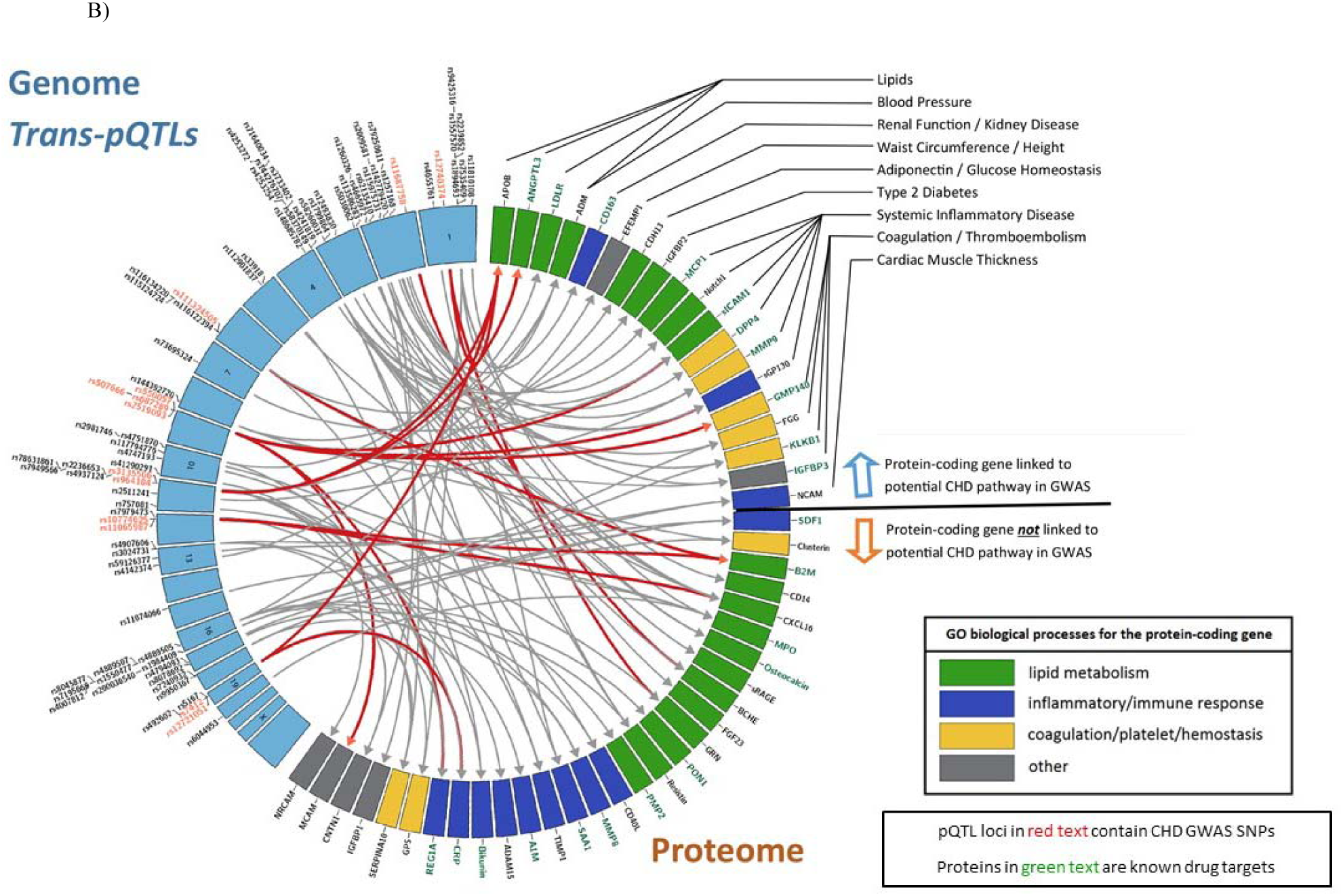
Sentinel *cis*- and *trans*-pQTLs and the Corresponding Proteins. Circos plots of sentinel *cis*- (Panel A) and *trans*-pQTLs (Panel B) and the plasma protein levels with which they are associated. Sentinel pQTLs are listed in order of chromosomal locations (blue boxes in the left semicircle). Loci containing pQTLs previously identified in GWAS to be associated with CHD are written in red text. Proteins with genome-wide significant pQTLs are listed in the right semicircle. The following three conditions are summarized for each protein: 1) The corresponding protein-coding gene is linked to a potential CHD pathway in previous GWAS (above the black line). 2) The corresponding protein-coding gene is a known drug target (green text). 3) GO biological processes for the protein-coding gene (green box denotes lipid metabolism pathways, blue box denotes inflammatory/immune response pathways, yellow box denotes coagulation/platelet/hemostasis pathways, and gray box denotes other pathways not included in the three most common, previously listed pathways). A single primary GO process was chosen when the protein-coding gene was included in multiple pathways. Abbreviations: CHD = coronary heart disease; GO = Gene Ontology; GWAS = genome-wide association study; pQTL = protein quantitative trait locus (i.e. genetic variant associated with protein level); SNP = single nucleotide polymorphism

The effect sizes and the proportion of inter-individual variation explained by some pQTLs were large. For example, *cis*-pQTL rs941590, is a missense variant that explained 32% of inter-individual variation in SERPINA10 levels (Figure S1) and was previously reported to be associated with family history of venous thrombosis.^18^ Three proteins (PON1, GRN, and LPA) had pQTLs that explained 10-30% of variation in protein levels. Minor allele frequency was inversely correlated with effect size, but not with proportion of variance explained. In general, *cis*-pQTLs and missense variants had larger effect sizes and explained a greater proportion of the variation in protein levels than did *trans*-pQTLs and non-coding variants, respectively (Figure 3).

**Figure 3.**
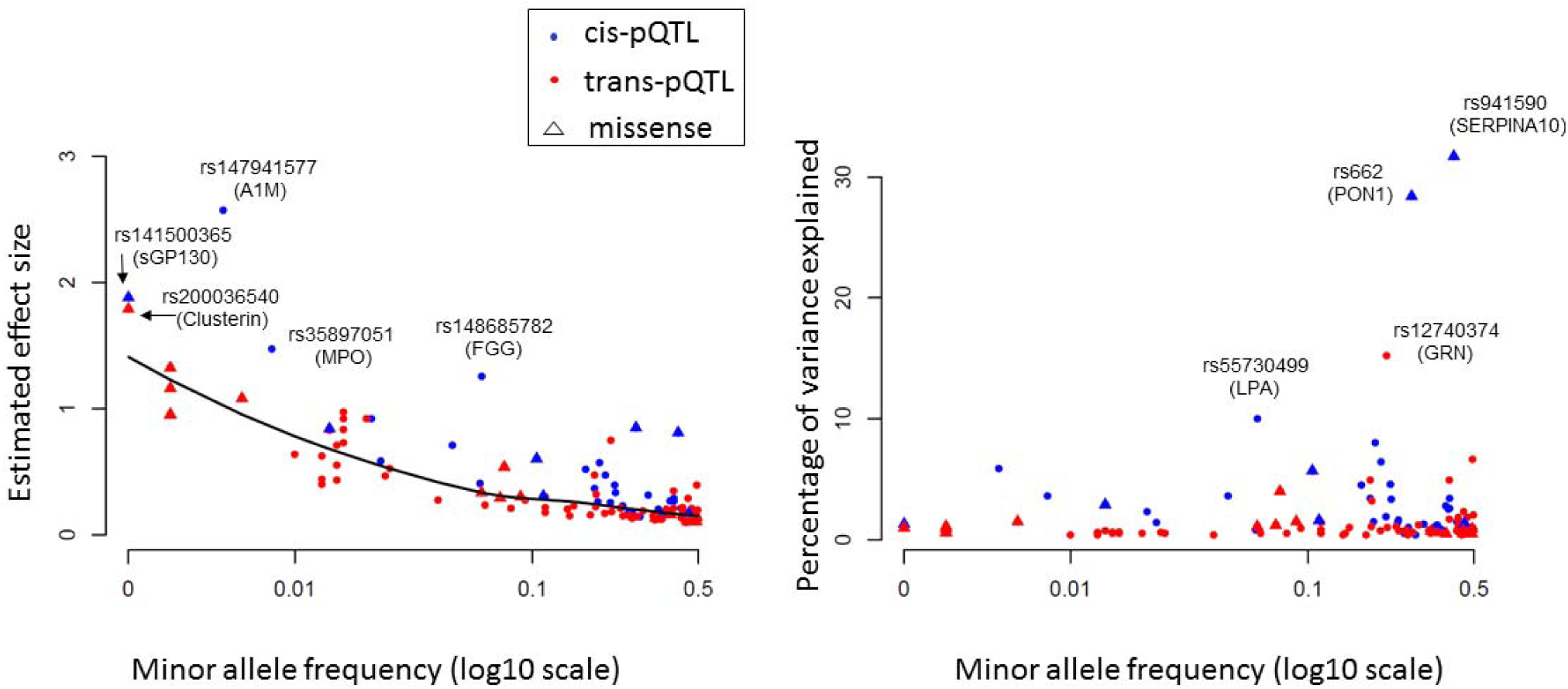
pQTL Minor Allele Frequency vs. Effect Size and Proportion of Variance Explained. Minor allele frequency of pQTLs (X-axis) vs. effect size of variants on proteins (left panel) and proportion of protein variance explained (right panel) for each sentinel pQTL locus. Abbreviations: pQTL = protein quantitative trait locus (i.e. genetic variant associated with protein level)

### External Replication

Among our 60 proteins linked to 130 sentinel pQTLs, 46 proteins (associated with 105 sentinel pQTLs) were measured in the KORA F4 study^12^ or in prior GWAS. For each pQTL locus identified in the FHS, the pQTL with the lowest protein-association p-value was selected as the sentinel pQTL for external replication. Thirty-six proteins, encompassing 82 sentinel pQTLs, were measured in KORA (21 *cis*-pQTLs and 61 *trans*-pQTLs); 23 additional loci were evaluated for replication in other GWAS. Based on 1,000 re-samplings of 1,000 unrelated FHS participants in the discovery sample, 34 pQTL-protein associations yielded p< 4.8x10^−4^ (alpha level of 0.05 after Bonferroni correction for 105 tests; 0.05/105) in ≥80% of samplings and thus were considered likely to replicate in a GWAS sample size of 1,000 (Table S9). Of the 21 qualifying sentinel *cis*-pQTLs from FHS, 10 replicated in KORA (p< 4.8x10^−4^; Table S10). Of the 61 sentinel *trans*-pQTLs (31 proteins) from FHS, 17 (14 proteins) replicated in KORA (p< 4.8x10^−4^; Table S10). Next, replication from other external protein GWAS was conducted. Overall, among sentinel pQTLs from FHS for which replication was possible, all of the ten most significant sentinel *cis*-pQTLs and seven of the ten most significant sentinel *trans*-pQTLs replicated. In total, 19 of 31 (61%) *cis*-pQTL loci and 22 of 74 (30%) *trans*-pQTL loci replicated at p< 4.8x10^−4^ (Table S10).

### pQTL Functional, Regulatory, and Clinical Annotation

Among the entire set of pQTLs, 334 are missense variants associated with 43 proteins and 8,217 are intronic variants (Table S11; Figure S2).^19^ Pathway enrichment analysis of all pQTLs using DEPICT^20^ identified 728 interrelated gene sets (false discovery rate [FDR]< 0.05, p< 0.0034); many are associated with lipids, metabolic processes, or inflammatory response (Table S12). Tissue enrichment analysis revealed that pQTL-mapped genes are highly expressed in monocytes (p=2.38x10^−4^) and hepatocytes (p=4.55x10^−5^). We employed Functional Mapping and Annotation^21^ of GWAS (FUMA; http://fuma.ctglab.nl) to generate detailed annotations of pQTLs for each protein (regional plot of each pQTL locus, functional categorization of pQTL SNPs, gene mapping, and pathway enrichment analyses) that are provided in Figure S3. These annotations revealed that pQTLs often reside in active regulatory regions and are frequently located in intergenic and intronic regions. Protein-specific Kyoto Encyclopedia of Genes and Genomes (KEGG) pathway analyses of the corresponding pQTLs revealed a preponderance of pathways concordant with the function of the studied protein.

### Enrichment of pQTLs with eQTLs

More than 90% pQTLs were annotated with chromatin marks or DNase hypersensitive sites by HaploReg^22^ (Table S13), suggesting that they play an important role in gene regulation. From the 1000G GWAS, we identified 8,542 pQTLs (46% of total discovery) for 20 proteins that also are whole blood eQTLs (genetic variants associated with gene expression levels in 5,257 FHS participants at FDR< 0.05),^23^ including 8,532 *cis*-eQTLs and 596 *trans*-eQTLs (Fisher’s exact test for enrichment p< 1x10^−8^; Table S14). Among the 130 sentinel pQTLs, 72 (55%) are eQTLs. Moreover, we identified pQTLs associated with expression of the corresponding protein-coding genes for 15 proteins, suggesting that many pQTLs affect circulating protein levels by regulating blood cell gene expression (Table S15).

### Clinical Annotation

We found that 58 missense pQTLs (from the Exome Chip) were linked to clinical disorders in the NCBI ClinVar^24^ database (Table S16). We identified examples where the missense pQTL and its associated protein are both linked to CHD-related traits. For example, rs2228671, a missense variant in the LDL-receptor gene (*LDLR*), was associated in our pQTL analysis with circulating APOB levels, the major lipoprotein of LDL particles, and was previously reported to be pathogenic for familial hypercholesterolemia (FH), a monogenic disorder of LDL cholesterol.^25^ Additionally, for several variants reported to be benign in ClinVar, we demonstrated associations with the disease-relevant protein, suggesting that they have clinical consequences.

### Integrating pQTLs with CHD-associated SNPs

We integrated our pQTLs with 2,738 CHD-related SNPs from the CARDIoGRAMplusC4D Consortium^1^ and other CHD GWAS.^2-4^ A total of 201 pQTLs (19 independent pQTLs with linkage disequilibrium r^2^< 0.2, representing 14 proteins) matched SNPs associated with CHD in GWAS (p< 5x10^−8^; Table S17). Table 1 displays the sentinel pQTL, based on lowest protein-association p-value that coincided with a CHD-related GWAS SNP, and the corresponding protein at each genetic locus. The proteins identified by this approach included GRN, APOB, ANGTL3, CRP, B2M, GMP140, sICAM1, REG1A, MCAM, LPA, sGP130, BCHE, sRAGE, and CXCL16.

**Table 1.** Proteins with pQTLs that Coincide with Coronary Heart Disease-associated SNPs from Genome-wide Association Studies.

| Protein | Protein-coding Gene | Location of Protein-coding Gene | pQTL* | pQTL Annotated Gene Locus | pQTL Location | pQTL Function | pQTL-protein P-value | CHD GWAS P-value† |
| --- | --- | --- | --- | --- | --- | --- | --- | --- |
| ANGPTL3 | *ANGPTL3* | 7:63.1M | rs964184 | *ZNF259* | 11:116.6Mb | UTR3 | $1.1 \times 10^{-14}$ | $8.0 \times 10^{-10}$ |
| APOB | *APOB* | 2:21.2M | rs12740374 | *CELSR2* | 1:109.8Mb | UTR3 | $3.2 \times 10^{-15}$ | $3.3 \times 10^{-18}$ |
| APOB | *APOB* | 2:21.2M | rs964184 | *ZNF259* | 11:116.6Mb | UTR3 | $9.5 \times 10^{-9}$ | $8.0 \times 10^{-10}$ |
| APOB | *APOB* | 2:21.2M | rs445925 | *APOC1* | 19:45.4Mb | Upstream | $7.0 \times 10^{-27}$ | $9.4 \times 10^{-11}$ |
| B2M | *B2M* | 15:45.0M | rs2508015 | *HLA-C* | 6:31.0Mb | Intergenic | $5.2 \times 10^{-10}$ | $1.5 \times 10^{-9}$ |
| B2M | *B2M* | 15:45.0M | rs10774625 | *ATXN2* | 12:119.1Mb | Intronic | $8.7 \times 10^{-11}$ | $8.0 \times 10^{-9}$ |
| BCHE | *BCHE* | 3:165.5M | rs35071165 | *ABI2* | 2:204.2Mb | Intronic | $3.4 \times 10^{-8}$ | $5.2 \times 10^{-10}$ |
| CRP | *CRP* | 1:159.7M | rs12721051 | *APOC1* | 19:45.4Mb | UTR3 | $2.4 \times 10^{-17}$ | $2.0 \times 10^{-10}$ |
| CXCL16 | *CXCL16* | 17:4.6M | rs11065987 | *CUX2* | 12:112.1Mb | Intergenic | $1.1 \times 10^{-8}$ | $2.5 \times 10^{-10}$ |
| GMP140 | *SELP* | 1:169.6M | rs2519093†† | *ABO* | 9:136.1Mb | Intronic | $1.2 \times 10^{-78}$ | $1.2 \times 10^{-11}$ |
| GRN | *GRN* | 17:42.4M | rs12740374 | *CELSR2* | 1:109.8Mb | UTR3 | $2.7 \times 10^{-268}$ | $4.6 \times 10^{-23}$ |
| LPA | *LPA* | 6:161.0M | rs55730499** | *LPA* | 6:161.0Mb | Intronic | $3.8 \times 10^{-167}$ | $5.4 \times 10^{-39}$ |
| MCAM | *MCAM* | 11:119.2M | rs550057 | *ABO* | 9:136.1Mb | Intronic | $1.4 \times 10^{-11}$ | $4.2 \times 10^{-9}$ |
| REG1A | *REG1A* | 2:79.3M | rs687289 | *ABO* | 9:136.1Mb | Intronic | $2.6 \times 10^{-12}$ | $7.7 \times 10^{-9}$ |
| sGP130 | *IL6ST* | 5:55.2M | rs507666†† | *ABO* | 9:136.1Mb | Intronic | $8.2 \times 10^{-19}$ | $1.6 \times 10^{-11}$ |
| sICAM1 | *ICAM1* | 19:10.4M | rs532436†† | *ABO* | 9:136.1Mb | Intronic | $7.7 \times 10^{-41}$ | $1.6 \times 10^{-11}$ |
| sRAGE | *AGER* | 6:32.1M | rs2523535** | *MICA* | 6:31.3Mb | Upstream | $1.3 \times 10^{-9}$ | $8.1 \times 10^{-10}$ |

\*For proteins with multiple pQTLs that coincided with coronary heart disease GWAS SNPs, the pQTL with the lowest p-value of association with its corresponding protein level is shown.
\*\*Indicates *cis*-pQTL. All other pQTLs shown in this table are *trans*-pQTLs.
†P-value of associations with coronary heart disease risk in GWAS reported in the NHGRI GWAS Catalog, GRASP Database, or CARDIOGRAMplusC4D Consortium.
††Indicates pQTLs located in the *ABO* gene that are in high LD ( $r^2 > 0.8$ ) with each other.
Abbreviations: CHD = coronary heart disease; GWAS = genome-wide association study; pQTL = protein quantitative trait locus (i.e. genetic variant associated with protein level)

We found the ABO locus to have links to CHD through five circulating proteins (MCAM, sICAM1, GMP140, sGP130, REG1A). ABO blood type has long been linked to CVD risk, including in the FHS,^26^ additional reports have linked the ABO locus to CVD via coagulation pathways.^27,28^ ABO locus-related proteins, identified in our study, are involved in inflammatory pathways, including interleukin and interferon signaling (Figure 2). The multi-protein association of this locus may be driven by the general function of ABO as a glycosyltransferase.

Some of the genes coding for CHD-related proteins have been linked to known CHD risk pathways in previous GWAS of lipids (APOB, LPA, ANGPTL3), coagulation (GMP140), and systemic inflammation (sGP130, sICAM1) (Figure 2). Many of the proteins that share genetic underpinnings with CHD are known drug targets (per the DrugBank database),^29^ or currently under development as such (*e.g.* ANGPTL3, LPA, sICAM1, GMP140). Several proteins with pQTLs linked to CHD, however, are not known drug targets, especially those from gene loci not previously linked to CHD risk pathways (e.g. BCHE, CXCL16, GRN, MCAM, and sRAGE).

### Causal Testing

We applied MR testing to infer the causal association between protein levels and CHD for all proteins having *cis*-pQTLs and those with at least four *cis*- or *trans*-pQTL loci that coincided with CHD-associated SNPs from GWAS.^1^ MR causally implicated LPA, REG1A, MCAM, and SAA1 via *cis*-pQTLs as instrument variables (p< 0.05; Table S18). For 11 proteins with pQTLs that coincided with SNPs from CHD GWAS and had at least four non-redundant *cis-* or *trans*-pQTLs, we conducted MR analyses using a multi-SNP approach, implemented in MRbase,^30^ which revealed causal CHD associations for APOB (p=0.0005) and GRN (p=7.0x10^−5^).

### Protein Associations with Clinical Outcomes

For 15 proteins (Figure 4) with pQTLs that coincided with CHD GWAS SNPs or tested positive in MR analyses at p< 0.05 we tested the associations of protein levels with a) major CHD (recognized myocardial infarction or CHD death; n=213 events) and b) CVD death (fatal CHD or death due to stroke, peripheral arterial disease, heart failure, or other CVD causes; n=199 events) occurring during a median follow-up of 14.3 years (25th percentile 11.4, 75th percentile 15.2 years) among 3,520 FHS participants age ≥50 years. Twelve of the 14 proteins with pQTLs that coincided with CHD GWAS SNPs were associated (nominal p< 0.05) with incident CHD or CVD death (Table 2). After adjusting for multiple testing of 15 proteins (p< 0.05/15 = p< 3.3x10^−3^), nine proteins remained associated with incident events. Four (REG1A, SAA1, APOB, and GRN) of the six proteins that tested causal for CHD by MR (at p< 0.05) were associated with CHD/CVD outcomes (at p< 0.05). Two proteins (AGP1 and HPX) that tested marginally positive for CHD risk in MR analysis (0.05< p< 0.10) were associated with new-onset CHD events (p=3.5x10^−8^ for AGP1 and p=2.0x10^−5^ for HPX). The protein effect sizes on CHD predicted from MR were consistent with the observed prospective protein-CHD associations (Figure 5).

**Figure 4.**
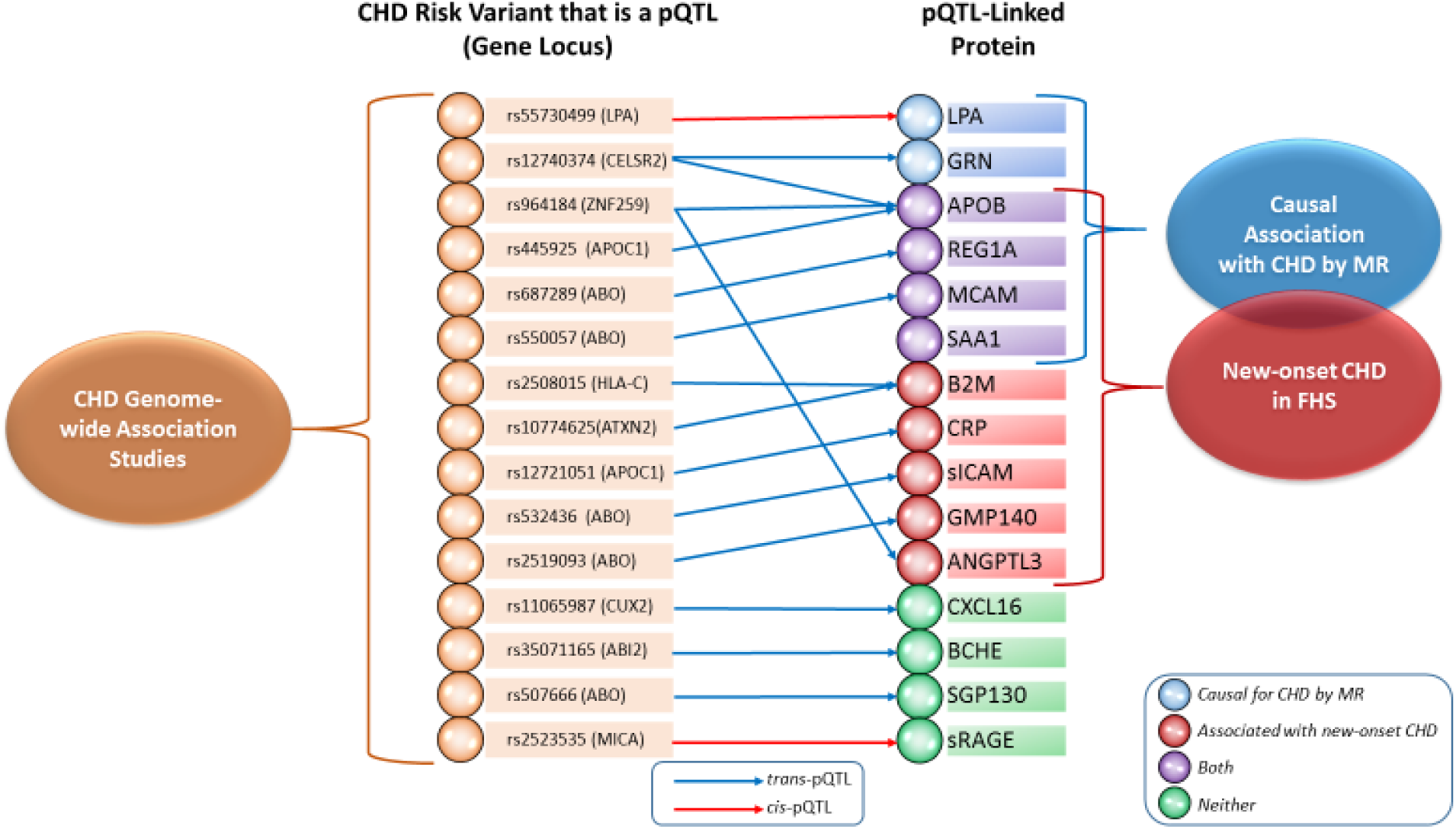
pQTL-Protein-Coronary Heart Disease Network. Network of proteins and significant pQTLs with annotated gene loci for the pQTLs that are also GWAS risk SNPs for CHD (see Table 1). For proteins with multiple pQTLs that coincide with coronary heart disease GWAS SNPs, the pQTL with the lowest p-value of association with its corresponding protein level is shown. The following two conditions are summarized: 1) Proteins that tested causal for CHD in Mendelian randomization (p< 0.05). 2) Proteins associated with new-onset CHD (p< 0.05) in 3,520 Framingham Heart Study participants age 50 years or older with long-term follow-up. Proteins in green fulfill neither condition 1 nor 2; proteins in blue fulfill condition 1; proteins in red fulfill condition 2; proteins in purple fulfill conditions 1 and 2. The pQTL rs2523535 for sRAGE was reported to be associated with CHD (p=8.1x10-10) in a Japanese GWAS (PMID 21971053). Abbreviations: CHD = coronary heart disease; FHS = Framingham Heart Study; MR = Mendelian randomization; pQTL = protein quantitative trait locus (i.e. genetic variant associated with protein level)

**Table 2.** Protein Associations with Coronary Heart Disease Events and Cardiovascular Disease Death in Framingham Heart Study Participants with Long-term Follow-up.

| Protein | Link to CHD in GWAS, MR, or Both | Protein Description | KEGG Pathways for Which pQTLs of Each Protein Are Enriched for† | Association with CHD Events* |  | Association with CVD Death* |  |
| --- | --- | --- | --- | --- | --- | --- | --- |
|  |  |  |  | Hazards Ratio (95% CI) | P-value†† | Hazards Ratio (95% CI) | P-value†† |
| ANGPTL3 | CHD GWAS | A member of a family of secreted proteins that function in angiogenesis | PPAR signaling pathway | 1.18 (1.01-1.36) | 0.03 | 0.99 (0.85-1.16) | 0.94 |
| APOB | Both | The main apolipoprotein of chylomicrons and low density lipoproteins | Endocytosis | 1.44 (1.24-1.67) | **1.8x10<sup>-6</sup>** | 1.07 (0.91-1.26) | 0.41 |
| B2M | CHD GWAS | A serum protein found in association with the major histocompatibility complex (MHC) class I heavy chain on the surface of nearly all nucleated cells | Type I diabetes mellitus<br>Antigen processing and presentation<br>Allograft rejection<br>Graft versus host disease<br>Autoimmune thyroid disease | 1.47 (1.24-1.75) | **9.0x10<sup>-6</sup>** | 1.97 (1.63-2.38) | **2.3x10<sup>-12</sup>** |
| BCHE | CHD GWAS | A cholinesterase enzyme and member of the type-B carboxylesterase/lipase family of proteins | None | 1.09 (0.94-1.26) | 0.3 | 0.82 (0.70-0.96) | 0.01 |
| CRP | CHD GWAS | A member of the pentaxin family | None | 1.4 (1.20-1.62) | **1.4x10<sup>-5</sup>** | 1.43 (1.23-1.68) | **5.6x10<sup>-6</sup>** |
| CXCL16 | CHD GWAS | A scavenger receptor on macrophages | Chemokine signaling pathway<br>Cytokine-cytokine receptor interaction | 1.13 (0.97-1.31) | 0.1 | 1.17 (1.00-1.37) | 0.04 |
|  |  |  | Intestinal immune network for IgA production<br>Neurotrophin signaling pathway |  |  |  |  |
| GMP140 | CHD GWAS | Ca(2+)-dependent receptor for myeloid cells that binds to carbohydrates on neutrophils and monocytes | Cell adhesion molecules | 1.23 (1.06-1.42) | $7.1 \times 10^{-3}$ | 1.25 (1.06-1.46) | $6.0 \times 10^{-3}$ |
| GRN | Both | A family of secreted, glycosylated peptides | None | 1.14 (0.98-1.32) | 0.08 | 1.29 (1.11-1.51) | **$1.2 \times 10^{-3}$** |
| LPA | Both | A serine proteinase that inhibits the activity of tissue-type plasminogen activator I | None | 1.09 (0.94-1.26) | 0.2 | 1.09 (0.93-1.27) | 0.29 |
| MCAM | Both | Plays a role in cell adhesion, and in cohesion of the endothelial monolayer at intercellular junctions in vascular tissue | None | 0.88 (0.76-1.03) | 0.1 | 1.11 (0.95-1.30) | 0.19 |
| REG1A | Both | A type I subclass member of the Reg gene family | Glycosphingolipid biosynthesis - lacto and neolacto series<br>Glycosphingolipid biosynthesis - globo series | 1.28 (1.10-1.48) | **$1.2 \times 10^{-3}$** | 1.47 (1.25-1.73) | **$2.6 \times 10^{-6}$** |
| SAA1 | MR | A member of the serum amyloid A family of apolipoproteins | Propanoate metabolism<br>Cysteine and methionine metabolism<br>Pyruvate metabolism | 1.28 (1.10-1.48) | **$1.0 \times 10^{-3}$** | 1.37 (1.17-1.59) | **$6.2 \times 10^{-5}$** |
| sGP130 | CHD GWAS | A signal transducer shared by many cytokines, including Interleukin 6 (IL6), ciliary neurotrophic factor (CNTF), leukemia inhibitory factor (LIF), and oncostatin M | None | 1.02 (0.88-1.18) | 0.8 | 1.34 (1.14-1.56) | **$2.7 \times 10^{-4}$** |
|  |  | (OSM) |  |  |  |  |  |
| sICAM1 | CHD<br>GWAS | A cell surface glycoprotein | Complement and<br>coagulation cascades | 1.23 (1.06-1.42) | $5.1 \times 10^{-3}$ | 1.33 (1.14-1.55) | **$2.1 \times 10^{-4}$** |
| sRAGE | CHD<br>GWAS | A member of the<br>immunoglobulin superfamily<br>of cell surface receptors | Antigen processing and<br>presentation<br>Type I diabetes mellitus<br>Allograft rejection<br>Graft versus host disease<br>Autoimmune thyroid<br>disease | 1.03 (0.89-1.20) | 0.7 | 1.18 (1.01-1.37) | 0.04 |

\*CHD events (n=213) included recognized myocardial infarction or death from CHD, and CVD death (n=199) included fatal CHD or death due to stroke, peripheral arterial disease, heart failure, or other CVD causes occurring during a median follow-up of 14.3 years (25th percentile 11.4, 75th percentile 15.2 years) among 3,520 Framingham Heart Study participants age $\geq 50$ years.
\*\*Proteins that tested positive in Mendelian randomization analysis for coronary heart disease risk at $0.05 < p < 0.10$ .
†For proteins with pQTLs that are enriched for more than five KEGG pathways, the top five most significant pathways based on enrichment p-value are shown.
††The p-value threshold for significance ( $p < 3.3 \times 10^{-3}$ ) was determined by the Bonferroni method ( $0.05/15$ proteins tested). Significant p-values are shown in **bold**.
Abbreviations: CHD = coronary heart disease; CI = confidence interval; CVD = cardiovascular disease; GWAS = genome-wide association study; KEGG = Kyoto Encyclopedia of Genes and Genomes; MR = Mendelian randomization

**Figure 5.**
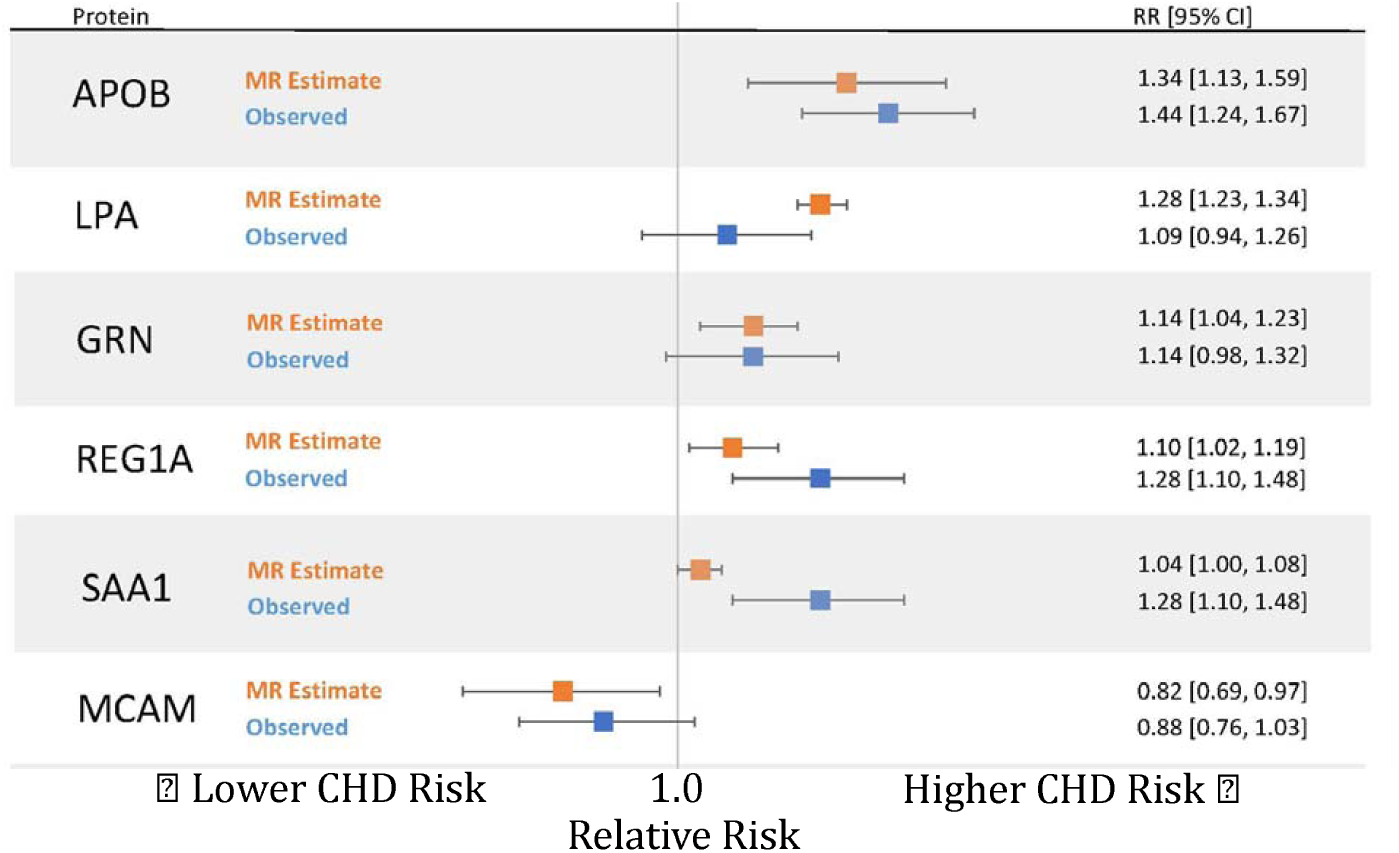
Comparison of Protein Effects on Coronary Heart Disease from Mendelian Randomization Estimate vs. Observed Hazards. Comparison of protein effects on CHD estimated from Mendelian randomization versus the observed hazards in 3,520 Framingham Heart Study participants with long-term follow-up. Abbreviations: CHD = coronary heart disease; CI = confidence interval; MR = Mendelian randomization; RR = relative risk

### Novel Proteins and Pathways Implicated in CHD

Whereas LPA and APOB can be viewed as positive controls because prior studies identified them as causal for CHD,^31-34^ four proteins that were causal for CHD in MR are novel, including REG1A, MCAM, SAA1, and GRN (Table S18). REG1A is a protein secreted by the pancreas and may be related to islet cell regeneration and diabetogenesis, potentially contributing to increased atherogenic risk of diabetes.^35^ REG1A levels were positively associated with CHD and CVD outcomes in our protein-trait association analyses (Table 2), and previous studies have shown that levels of REG1A are elevated in individuals with CHD and type 2 diabetes.^35,36^ MCAM, also known as CD146, a trans-membrane glycoprotein, is highly expressed in vascular cells and plays a role in cell adhesion. MCAM/CD146 levels are a biomarker of endothelial activation/injury and are associated with carotid intima thickness^37,38^ and risk for acute CHD events.^39-41^ Our MR and integrative GWAS analyses suggest a protective role of MCAM/CD146 on CHD risk. Protein-trait association analysis for MCAM/CD146 in the FHS did not find evidence of association with CHD or CVD events, however, a previous FHS case-control analysis reported an inverse association of MCAM/CD146 with myocardial infarction,^36^ concordant with our MR results.

SAA1, a precursor to amyloid A, is an inflammatory apolipoprotein inversely associated with HDL-cholesterol.^42^ Serum SAA1 levels are elevated in patients with CHD.^43^ In addition, SNPs located in the *SAA1* gene are associated with carotid intima media thickness and HDL-cholesterol levels.^44,45^ Our follow-up analyses revealed association of circulation SAA1 levels with CHD and CVD risk. MR results for GRN were positive, largely by virtue of sentinel *trans*-pQTLs. One of our sentinel *trans*- pQTLs for GRN, rs12740374, is located at the *CELSR2/SORT1* locus on Chromosome 1; it explained 15% of variance in GRN levels and was associated with CHD at p< 4.6x10^−23^ in prior GWAS.^1^ Previous studies reported that rs12740374 affects expression levels of the *SORT1* gene in human hepatocytes, which in turn regulate LDL-cholesterol levels.^46,47^ Our longitudinal analyses revealed association of GRN with CVD death in FHS participant (p=0.001; Table 2).

### Molecular QTL browser

Our pQTL resource is accessible through the NCBI Molecular QTL Browser (ftp://ftp.ncbi.nlm.nih.gov/eqtl/original_submissions/FHS_pQTLs/; a link to the browser will be sent to reviewers under separate cover), which serves as a data resource for associations between genetic variants and molecular phenotypes. The browser links our pQTL results to eQTLs and other molecular resources via a user-friendly interface. Users can browse and search results and specify p-value cutoffs and other data filters. The Molecular QTL Browser also permits users to conduct targeted studies of specific genes based on prior evidence. The integrated data resource enables searches across datasets and filtering by functional annotation and genomic position.

## Discussion

Using a multistage strategy, we discovered thousands of pQTLs associated with scores of proteins that were selected *a priori* as high-value plasma proteins for CVD. Integration of pQTLs with CHD GWAS revealed 14 proteins with pQTLs that coincide with CHD SNPs (Table 1) and MR analyses identified six proteins with evidence for causal effects on CHD (Table S18), four of which were novel (REG1A, MCAM, SAA1, and GRN). Furthermore, most of these proteins were associated with new-onset CHD or CVD events in FHS participants with long-term follow-up (Table 2). Our strategy connected pQTLs, genes, the proteins they code, and CHD risk (Figure 4) and highlighted a comprehensive approach to bridge the GWAS knowledge gap for genetic variants that have no links to disease via known mechanisms.

We acknowledge several limitations of our study. Participants were of European ancestry; consequently, the results may not be directly generalizable to populations with different genetic backgrounds. Although our sample size for GWAS was large, our ability to detect pQTLs and to test them for causality using MR was limited by power. Protein levels were measured in whole blood and may not reflect tissue-specific patterns of expression.

To our knowledge, this is the largest sample size pQTL study with well-powered discovery, independent external replication, CHD causal testing, and confirmatory prospective protein-CHD outcome findings. We provide a large and comprehensive compilation of pQTLs as a resource for other researchers (via the NCBI Molecular QTL Browser) and provide evidence that an integrated genomic approach can identify proteins with putatively causal effects on CHD. Although some of our causally-implicated proteins may act through classic CHD risk factors and known pathways, many do not and thus represent attractive candidate targets for drug development. Additional studies are needed to elucidate the mechanisms by which such proteins alter CHD risk as well as trials to confirm our MR prediction that perturbing these pathways can prevent CHD events. Taken together, the genetic variants associated with circulating protein levels in this study shed new light on genes, proteins, and pathways contributing to the pathogenesis of CHD, which could have profound implications for the treatment and prevention of the leading cause of death worldwide.

## Acknowledgements

Framingham Heart Study: The Framingham Heart Study is funded by National Institutes of Health contract N01-HC-25195. This project was funded in part by the Division of Intramural Research, National Heart, Lung, and Blood Institute (NHLBI), National Institutes of Health (NIH), Bethesda, MD. The views expressed in this manuscript are those of the authors and do not necessarily represent the views of the National Heart, Lung, and Blood Institute; the National Institutes of Health; or the U.S. Department of Health and Human Services. Dr. Ho is supported in part by NIH grant K23-HL116780 and a Massachusetts General Hospital Hassenfeld Research Scholar Award. We thank all the study participants who helped to create this valuable resource and supported this work. We thank the data management group of FHS for organizing and providing these data. We thank the National Institutes of Health Fellows Editorial Board members for their valuable edits and comments. This study used the high-performance computational capabilities of the Biowulf Linux cluster at the National Institutes of Health, Bethesda, MD.

KORA: KS was supported by ‘Biomedical Research Program’ funds at Weill Cornell Medicine in Qatar, a program funded by the Qatar Foundation. The KORA study was initiated and financed by the Helmholtz Zentrum München – German Research Center for Environmental Health, which is funded by the German Federal Ministry of Education and Research (BMBF) and by the State of Bavaria. Furthermore, KORA research was supported within the Munich Center of Health Sciences (MC-Health), Ludwig-Maximilians-Universität, as part of LMUinnovativ. The KORA-Study Group consists of A. Peters (speaker), J. Heinrich, R. Holle, R. Leidl, C. Meisinger, K. Strauch, and their co-workers, who are responsible for the design and conduct of the KORA studies. We gratefully acknowledge the contribution of all members of field staff conducting the KORA F4 study. Most of all, we thank all study participants for their invaluable contributions to this study.

## Methods

### Study Design

The study consisted of seven steps (Figure 1): 1) selection and measurement of 71 high-value plasma proteins for atherosclerotic CVD via multiplex immunoassays in 7,333 FHS participants, 2) genome-wide association study of the 71 proteins in 6,861 FHS participants to identify genome-wide significant pQTLs, 3) functional enrichment analyses of the identified pQTLs, 4) independent external replication of the sentinel pQTLs in KORA, 5) integrative analysis to pQTLs that coincide with CHD SNPs from GWAS, 6) identification of causal proteins for CHD using a Mendelian randomization approach, 7) association analysis of proteins from steps 5 and 6 with risk for incident CHD death and CVD death in 3,520 FHS participants age 50 years or older with available long-term follow-up.

### Discovery Study Sample

The FHS is a community-based prospective study of CVD and its risk factors that recruited three generations of participants within families in 1948, 1971, and 2002, respectively.^48-50^ The study samples for this investigation were collected from 7,333 participants from the FHS Offspring (Exam 7; 1998-2001) and Third Generation (Exam 1; 2002-2005) cohorts. The final sample for GWAS was composed of 6,861 participants with complete imputed dosage data based on the 1000 Genomes Project reference panel (1000G).^17^ For association analyses using Exome Chip genotypes (see Genotyping for details), the sample size was 6,763. Genome-wide analysis of SNPs associated with gene expression levels (eQTLs) was performed on 5,257 FHS participants in whom both genotype and gene expression data were available.^23^

### Replication Study Sample

The KORA F4 study is a prospective population-based cohort study consisting of 3,080 participants living in Augsburg, Southern Germany.^12,51^ A total of 1,000 participants who also participated in a metabolomic study with follow-up information for aging-related diseases composed the study population for replication. After excluding participants with missing genotype or protein data (n=3), the final KORA sample included 997 individuals.

### Power Calculation

For power in the discovery stage with n=6,800, we assumed an additive genetic model with no interaction and a population mean=0 and standard deviation=1 for all rank-normalized protein levels. At α=5x10 for a two-sided test, power was estimated for MAF=0.002, 0.005, 0.01, 0.05, 0.1, 0.2, 0.3, 0.4, and 0.5 with QUANTO.^52^ For empirical power in the replication stage, we performed pQTL analysis with 1,000 resamplings of 1,000 unrelated FHS participants. We counted the number of tests with p< 0.05/n in the 1,000 resamplings, where n is the number of pQTLs that tested for replication in KORA.

### Clinical Measures

All FHS participants underwent periodic clinical examinations with standard protocols as described previously.^50^ A three-physician panel was formed to perform medical chart review weekly. The review panel jointly assigned CVD diagnoses and causes of death. All suspected CVD events were adjudicated by the physician-panel after reviewing all available medical evidence including hospital records, personal physician records, and interviews with next of kin in the event of an out-of-hospital death. Recognized myocardial infarction (MI) was diagnosed when two of three of the following conditions were present: prolonged chest discomfort or symptoms of coronary ischemia, elevated biomarkers of myocardial necrosis (e.g. CK-MB or troponin), and the development of new diagnostic Q-waves on the ECG. Fatal CHD events included fatal MI and other deaths due to CHD as an underlying cause in the absence of evidence of recent infarction. Fatal CVD events additionally included deaths due to stroke, peripheral arterial disease, heart failure, or other cardiovascular causes.

### Protein Quantification

FHS fasting blood plasma samples were collected and stored at - 80°C. Candidate protein biomarkers were selected *a priori* based on previous evidence of association with atherosclerotic CVD or its risk factors using the following complementary approaches: a) comprehensive literature search,^53^ b) proteomics discovery via mass spectrometry in the FHS or elsewhere,^36,54^ and c) targeting proteins coded by genes identified via gene expression profiling studies^55,56^ or GWAS^57^ of atherosclerotic CVD and its risk factors. A total of 85 plasma protein biomarkers were assayed using a modified enzyme-linked immunosorbent assay sandwich method, multiplexed on a Luminex xMAP platform (Luminex, Inc., Austin, TX). All targets were first developed as singleton assays before compatible targets were pooled to create multiplex panels. Standard Luminex assays with previously published methods were used.^58,59^ Measurements were calibrated using a seven-point calibration curve (in triplicate) and tested for recovery at both ends of the quantitation scale. The ‘High’ and ‘Low’ spike controls (QC1 and QC2 respectively) were used to calculate intra- and inter-assay coefficients of variation (CV) for each protein. A total of 14 proteins had low call-rate (< 90%) mainly due to values falling below the lower detection limit that were excluded for the current study. A list of the 71 proteins and their coefficients of variation and selection criteria were shown in Table S1.

For the KORA study, plasma levels of 1,129 proteins in 1,000 blood samples were measured using the SOMAscan platform (SomaLogic Inc., Boulder, Colorado), a multiplexed aptamer-based affinity proteomics platform; 1,124 proteins passed quality control. Protein measurement protocol, normalization of protein values, and data quality are described elsewhere.^12^

### Genotyping

Genotyping and QC methods in the FHS have previously been described.^17^ In brief, genome-wide genotyping was conducted using the Affymetrix 500K mapping arrays and 50K supplemental Human Gene Focused arrays (Affymetrix, Inc., Santa Clara, CA) as well as the Illumina Human Exome BeadChip v.1.0 (Exome Chip; Illumina, Inc., San Diego, CA). Genotypes from the Affymetrix arrays were used in conjunction with the 1000G reference panel^17^ to generate an imputed set of ∼30 million variants using MACH.^60^ SNPs with imputation quality ratio < 0.3 (imputation quality ratio is calculated by the ratio of the variances of the observed and the estimated allele counts) or minor allele frequency (MAF) < 0.01 were excluded, leaving a final set of 8,509,364 SNPs for 1000 genomes imputed GWAS.

The Exome Chip includes rare coding variants not covered by previous genotyping arrays.^61^ More than 90% of the SNPs included in the Exome Chip are non-synonymous variants, splice variants, or stop codon altering variants. Common variants on the Exome Chip include 5,542 SNPs that were selected based on their associations with disease traits reported in the NHGRI GWAS Catalog.^1^ Rare variants with MAF< 1x10^−4^ were excluded from analysis.

For KORA, the Affymetrix Axiom Array (Affymetrix, Inc., Santa Clara, CA) was used to genotype 3,788 study participants.^12,51^ Genotypes were then imputed from the 1,000G reference panel^15^ and used for lookup of the replication targets.

### Functional Annotation of pQTLs

We used HaploReg^22^ v4.1 to functionally annotate our pQTL results. Using information from the Roadmap Epigenomics^62^ and ENCODE projects,^63^ HaploReg linked SNPs and small insertions/deletions with chromatin state, protein binding annotation, and regulatory motifs. A total of 14,756 pQTLs could be found in the HaploReg database. We used DEPICT^20^ to conduct gene prioritization, pathway analysis, and tissue/cell type enrichment analysis. DEPICT used information from co-regulation of gene expression from 77,840 samples, in conjunction with 14,461 reconstituted functional gene sets, to assess pathway enrichment and prioritize genes. In addition, DEPICT utilized a set of 37,427 human microarrays to identify enrichment of highly expressed genes in specific tissue/cell types. We used Functional Mapping and Annotation^21^ of GWAS (FUMA; http://fuma.ctglab.nl) to categorize proteins based on known pathways and conduct functional annotation of pQTLs (regional plot of each pQTL locus, functional categorization of pQTL SNPs, gene mapping, and pathway enrichment analyses).

### Gene Expression

Gene expression profiling was conducted using the Affymetrix Human Exon 1.0 ST GeneChip platform (Affymetrix Inc., Santa Clara, CA), comprised of > 5.5 million probes covering expression of 17,873 mRNA transcripts. Gene expression values were normalized and adjusted for three technical covariates (batch, first principal component, and residual of probeset mean values) as described previously.^23^

### Coronary Heart Disease-associated SNPs

The CARDIoGRAMplusC4D Consortium^1^ GWAS of CHD yielded 1,892 genome-wide significant SNPs (at p< 5x10^−8^) from 1000G imputation. The National Human Genome Research Institute (NHGRI) GWAS catalog^2^ (downloaded in July 2016) and Genome-wide Repository of Associations Between SNPs and Phenotypes (GRASP)^3,4^ v.2.0 (downloaded in June 2016) included 846 SNPs associated in GWAS with CHD at genome-wide significance level (p< 5x10^−8^).

### Statistical Methods

Statistical analyses in the FHS were performed using R software version 3.1.1^64^ or SAS software version 9.4.

### Genome-wide association (pQTL) analyses

Linear mixed effects models (the “LMEKIN” function of Kinship Package in R^64^) were used to test associations of inverse-rank normalized protein levels with 1000G or Exome Chip variants in conjunction with an additive genetic model. We applied a p-value threshold of 5x10^−8^ for defining significant pQTLs. A *cis*-pQTL was defined as a SNP residing within 1 megabase (Mb) upstream or downstream of the transcription start site of the corresponding protein-coding gene. A SNP located > 1 Mb upstream or downstream of the gene transcript or on a different chromosome from its associated gene was categorized as a *trans*-pQTL.

Linkage disequilibrium (LD) was computed as the square of Pearson’s correlation (r^2^) between imputed additive dosages of genotypic variants within the same chromosome across 8,481 FHS individuals with genotype data. Independent pQTLs for a given protein were defined as those with LD r^2^< 0.2 with other pQTLs at a genomic locus. For a genetic locus with multiple pQTLs in LD (*i.e.,* LD r^2^> 0.2), we selected the pQTL with the lowest p-value to represent the sentinel pQTL for that locus.

For KORA, linear regression models were performed on the follow-up SNPs using R version 3.1.3.^64^ Associations between inverse-normalized protein levels and imputed dosages were tested using linear additive genetic regression models adjusted for age, sex, and body mass index.^12^

### eQTL Mapping

We used linear mixed effects models, accounting for familial relationships using “PEDIGREEMM” in R,^64^ to assess associations between ∼8.5 million 1000G SNPs that were additively coded and expression levels of 17,873 transcripts.^23^

Models were adjusted for age, sex, platelet count, differential white cell count (percentages of lymphocyte, monocyte, eosinophil, and basophil), and for 20 PEER factors^65,66^ to reduce confounding due to unmeasured factors. The criteria used to define *cis* and *trans* effects for pQTLs were also applied to eQTLs. A false discovery rate (FDR) threshold of 0.05 was applied separately for *cis*- and *trans*-eQTLs.

### Mendelian randomization

We used an MR approach to test for causal associations between protein biomarkers and CHD risk. The sentinel *cis*-pQTL for each protein, based on lowest p-value of association in either 1000G GWAS or Exome Chip analysis, was selected as the instrumental variable (IV) for its perspective protein in MR analysis. Based on the association between the sentinel *cis*-pQTL and CHD in prior GWAS,^1^ a putative causal effect of one standard error difference in inverse-rank normalized protein level on CHD was calculated as the per risk allele effect on CHD risk dependent on the per risk allele effect on one standard error difference in inverse-rank normalized protein level (the Wald ratio test).^67^ For proteins with suggestive single *cis*-pQTL results (0.05< p< 0.1) and with additional non-redundant *cis*-pQTLs, we conducted single-locus multi-SNP MR. Low-level correlation (LD r^2^< 0.2) between variants in the genetic risk score was adjusted for in MR analysis using the method developed by Burgess et al.^68^ Similarly, for proteins with pQTLs that shared genetic signals with CHD from GWAS, we conducted multi-SNP MR using MRbase^30^ when there were at least four non-redundant pQTL loci.

### Associations of protein levels with CVD

To analyze associations between plasma protein levels and MI/CHD death and CVD death in FHS participants, protein biomarkers were rank-normalized. Cox proportional hazard models were used to predict MI/CHD death and CVD death for each biomarker, adjusting for age and sex. Participants younger than 50 years of age at baseline were excluded from outcome analyses due to a paucity of events in this age group. In addition, participants with prevalent MI/CHD or CVD at baseline were excluded from analyses of incident events, leaving a final sample size of 3,520 FHS participants.

### Independent External Replication

After merging our 1000G and Exome Chip GWAS results, the pQTL with the lowest p-value of association at each genetic locus was selected as the sentinel pQTL. We conducted independent external replication of our sentinel pQTLs in the KORA study^12^ and in other protein GWAS. Out of the 60 proteins with pQTL SNPs in the FHS, replication was conducted for 47 proteins from discovery that also were measured in KORA or other studies. The sentinel pQTL at each genetic locus in the FHS was determined to be successfully validated if its corresponding 1000G-imputed genotype or strong proxy (LD r^2^> 0.8) in KORA was also a significant pQTL for the corresponding protein and if directionality of pQTL-protein association was preserved. Statistical significance was defined as a p-value < 0.05/n (n was the number of pQTLs that were studied in KORA).

### Study Approval

All participants from the FHS and KORA study gave informed consent for participation in this study and for the collection of plasma and DNA for analysis. The KORA study was approved by the Ethics Committee of the Bavarian Medical Association, Germany.

### Data Access

All data from the FHS for this study are accessible (dbGaP Study Accession: phs000363.v16.p10). Data for KORA are available upon request from KORA-gen (http://epi.helmholtz-muenchen.de/kora-gen). Requests are submitted online and are subject to approval by the KORA board.

